# Metabolomic profiling revels systemic signatures of premature aging induced by Hutchinson-Gilford Progeria Syndrome

**DOI:** 10.1101/554220

**Authors:** Gustavo Monnerat, Geisa Paulino Caprini Evaristo, Joseph Albert Medeiros Evaristo, Caleb Guedes Miranda dos Santos, Gabriel Carneiro, Leonardo Maciel, Vânia Oliveira Carvalho, Fábio César Sousa Nogueira, Gilberto Barbosa Domont, Antonio Carlos Campos de Carvalho

**Affiliations:** Institute of Biophysics Carlos Chagas Filho, Federal University of Rio de Janeiro, Rio de Janeiro, Brazil; Laboratory of Proteomics, LADETEC, Institute of Chemistry, Federal University of Rio de Janeiro, Rio de Janeiro, Brazil; Instituto de Biologia do Exército, Ministério da Defesa do Brasil; Department of Pediatrics, Federal University of Paraná, Curitiba, Brazil; Proteomics Unit, Institute of Chemistry, Federal University of Rio de Janeiro, Rio de Janeiro, Brazil; National Institute of Cardiology, Rio de Janeiro, Brazil

**Keywords:** Lamin, HGPS, Premature aging, Metabolomics, Biomarkers, Meta-bolic profiling

## Abstract

Hutchinson-Gilford Progeria Syndrome (HGPS) is an extremely rare genetic disorder. HGPS children present a high incidence of cardiovascular complications along with altered metabolic processes and accelerated aging process. No metabolic biomarker is known and the mechanisms underlying premature aging are not fully understood. The present study analysed plasma from six HGPS patients of both sexes (7.7±1.4 years old; mean±SD) and eight controls (8.6±2.3 years old) by LC-MS/MS in high-resolution non-targeted metabolomics (Q-Exactive Plus). Several endogenous metabolites with statistical difference were found. Multivariate statistics analysis showed a clear separation between groups. Potential novel metabolic biomarkers are identified using the multivariate area under ROC curve (AUROC) based analysis, showing an AUC value higher than 0.80 using only two metabolites, and reaching 1.00 when increasing the number of metabolites in the AUROC model. Targeted metabolomics was used to validate some of the metabolites identified by the non-targeted method. Taken together, changed metabolic pathways in that panel involve sphingolipid, amino acid, and oxidation of fatty acids among others. In conclusion our data show significant alterations in cellular energy use and availability, in signal transduction, and in lipid metabolites, creating new insights on metabolic alterations associated with premature aging.

## Introduction

Hutchinson-Gilford Progeria Syndrome (HGPS) is an extremely rare genetic disorder. Children with HGPS present a high incidence of severe cardio-vascular complications along with altered metabolic processes, associated with an accelerated aging process (*1-4*). Despite a great increase in the scientific knowledge about HGPS, no specific biomarker is known for HGPS and the underlying molecular mechanisms are not fully understood. HGPS is induced by a single mutation in the LMNA gene, creating a mutant protein isoform with deletion of 50 amino-acids near in the protein Lamin A. The mutated protein, known as progerin (isoform 6), is toxically accumulated in the cells. Progerin, despite being able to enter the cell nucleus, does not incorporate normally into the nuclear membrane lamina, leading to several abnormalities in nuclear trafficking(*5, 6*). Interestingly, unaffected aged individuals show a similar splice event, leading to progerin expression that may play a role in cellular senescence(*6*).

Aging is the biological process of gradually accumulating deleterious changes in cells, decreasing the physiological capacity(*7, 8*). Aging is not considered a disease, but it intensely rises the risk of developing chronic cardio-vascular(*9*) and metabolic diseases(*10*). It is known that metabolic systemic profiles are age-dependent, reflecting metabolism alterations, such as incomplete fatty acid mitochondrial oxidation(*11-13*).

Metabolomics is, among other “omics” strategies, one of the most complete and reliable sources of information for circulatory mediator analysis, biomarker discovery pipeline and mechanistic disease investigation(*14*). In the present study, we applied metabolomics to samples from HGPS patients and identified several metabolites from different biological pathways dysregulated. Multivariate and univariate statistical analysis demonstrated metabolic pathways and potential new biomarkers that might act as central mediators in this syndrome and in senescence.

## Results

We used non-targeted based metabolomics to investigate metabolites differentially expressed in plasma samples obtained from 6 HGPS patients and 8 healthy donors, as summarized in Table S1, aiming to identify new biomarkers and novel mechanisms of the disease.

In order to avoid the inclusion of exogenous compounds in our analysis, contaminants, medications, and their metabolites, food and flavouring compounds were excluded from the metabolite list, resulting in a final feature list of 40 known molecules of endogenous origin presenting the statistical difference between the two groups. Each of the identified metabolites was found to have false discovery rates (FDRs) of less than 10%. Information regarding the metabolites identified in the present study is available in Table 1. Figure S1 shows typical total extracted ion chromatograms of all analyzed samples, demonstrating the efficient separation of the plasma compounds and reproducibility. The deuterated internal standard spiked in during sample preparation was used to calculate the coefficient of variation (CV) of our method. Figure S2 demonstrates that our CVs were <15% among samples.

**Table 1.** Significant dysregulated metabolites of HGPS

| Metabolite | Class | HMDB ID | Fold Change | p-value | FDR |
| --- | --- | --- | --- | --- | --- |
| LysoPC(17:0) | Glycerophospholipid | HMDB0012108 | 0.57 | 0.0179 | 0.12 |
| LysoPC(16:0) |  | HMDB0010382 | 0.50 | 0.0007 | 0.02 |
| PC(16:0/18:2) |  | HMDB0011211 | 0.45 | 0.0004 | 0.01 |
| Glucose | Carbohydrates and carbohydrate conjugates | HMDB0000122 | 3.66 | 0.0001 | 0.01 |
| N(6)-Methyllysine | Carboxylic acids and derivatives | HMDB0002038 | 7.74 | 0.0238 | 0.13 |
| Arginine |  | HMDB0000517 | 2.48 | 0.0056 | 0.06 |
| gamma-Aminobutyric acid |  | HMDB0000112 | 1.80 | 0.0413 | 0.19 |
| Aminolevulinic acid |  | HMDB0001149 | 0.77 | 0.0331 | 0.17 |
| Phenylalanine |  | HMDB0000159 | 0.54 | 0.0003 | 0.01 |
| N-Phenylacetylglutamine |  | HMDB06344 | 0.45 | 0.0172 | 0.12 |
| Pyroglutamylglycine |  | HMDB0061890 | 0.32 | 0.0016 | 0.02 |
| Aceglutamide |  | HMDB0006029 | 0.29 | 0.0011 | 0.02 |
| Threoninyl-Aspartate |  | HMDB0029057 | 0.27 | 0.0005 | 0.01 |
| Oleoylcarnitine | Fatty Acyl | HMDB0005065 | 0.44 | 0.0071 | 0.07 |
| L-Carnitine |  | HMDB0000062 | 1.72 | 0.0087 | 0.08 |
| Acetyl-L-carnitine |  | HMDB0000201 | 11.23 | 0.0312 | 0.16 |
| 3-Dehydroxycarnitine |  | HMDB0006831 | 0.75 | 0.0275 | 0.15 |
| Palmitoylcarnitine |  | HMDB0000222 | 0.64 | 0.0263 | 0.15 |
| Decanoylcarnitine |  | HMDB0000651 | 0.47 | 0.0411 | 0.19 |
| 9-Decenoylcarnitine |  | HMDB0013205 | 0.44 | 0.0005 | 0.01 |
| Phosphatidylcholine(16:0/16:0) | Glycerophospholipids | HMDB0011206 | 0.67 | 0.0015 | 0.02 |
| 1-octadecylglycero-3-phosphocholine |  | HMDB0062195 | 0.57 | 0.0066 | 0.07 |
| Glycerylphosphorylcholine |  | HMDB0000086 | 0.51 | 0.0049 | 0.06 |
| Dipalmitoylphosphatidylcholine |  | HMDB0000564 | 0.47 | 0.0029 | 0.04 |
| Lactide | Hydroxy acids and derivatives | not found | 3.62 | 0.0001 | 0.01 |
| 5-Hydroxytryptophol | Indoles and derivatives | HMDB0001855 | 6.65 | 0.0121 | 0.10 |
| 4,6-Dioxoheptanoic acid | Keto acids and derivatives | HMDB00635 | 2.04 | 0.0032 | 0.04 |
| LysoPC(P-16:0) | Lysophospholipid | HMDB0010407 | 0.35 | 0.0003 | 0.01 |
| Choline | Organonitrogen compounds | HMDB0000097 | 0.57 | 0.0002 | 0.01 |
| PC(o-18:1(11Z)/16:0) | Phosphatidylcholine | HMDB0013424 | 0.77 | 0.0342 | 0.17 |
| PC(O-16:0/18:2(9Z,12Z)) |  | HMDB0011151 | 0.51 | 0.0074 | 0.07 |
| N-Methylethanolaminium phosphate | Phosphoethanolamine | HMDB0060173 | 0.46 | 0.0367 | 0.18 |
| (3beta,19alpha)-3,19,23,24-Tetrahydroxy-12-oleanen-28-oic acid | Prenol lipids | HMDB0040784 | 0.23 | 0.0168 | 0.12 |
| Pyridoxamine | Pyridines and derivatives | HMDB0001431 | 0.56 | 0.0142 | 0.11 |
| Citicoline | Pyrimidine nucleotides | HMDB0001413 | 0.55 | 0.0055 | 0.06 |
| 13-cis retinol | Retinoids | HMDB0006221 | 2.16 | 0.0156 | 0.12 |
| Palmitoyl sphingomyelin | Sphingolipids | HMDB0061712 | 0.70 | 0.0136 | 0.11 |
| Pregnenolone | Steroids and steroid derivatives | HMDB0000253 | 2.10 | 0.0188 | 0.12 |
| (3beta,24R,24'R)-fucosterol epoxide | Sterol Lipids | not found | 0.43 | 0.0389 | 0.19 |
| Bilirubin | Tetrapyrroles and derivatives | HMDB0000054 | 6.34 | 0.0128 | 0.10 |

A data matrix including the average area values of the uniquely identified analyzed compounds in each sample was generated. Multivariate statistics using both unsupervised and supervised strategies were then applied to the data. Unsupervised PCA of the metabolomics data demonstrated a clear separation between groups (Figure 1a). Percent Variance Captured by PCA Model for the Principal Component 1 (PC1) was 55.4%, and for Principal component 2 (PC2) was 6.9%. The green (patients) and red (controls) areas in Figure 1a represent the 95% confidence intervals for each group. The application of supervised PLS-DA also permits a detailed group separation between HGPS and control cohorts as shown in Figure 1b. The PLS-DA model captured 55.4% of the variance in component 1 and 5.1% in component 2. The components of the PLS-DA models were used to predict the accuracy (Accuracy) based on the cross-validation, the sum of squares captured by the model (R2), and the cross-validated R2 (Q2). The PLS-DA cross-validation data are summarized together with a set of permutation tests demonstrating statistical significance in the PLS model (Figure S3).

**Figure 1.**
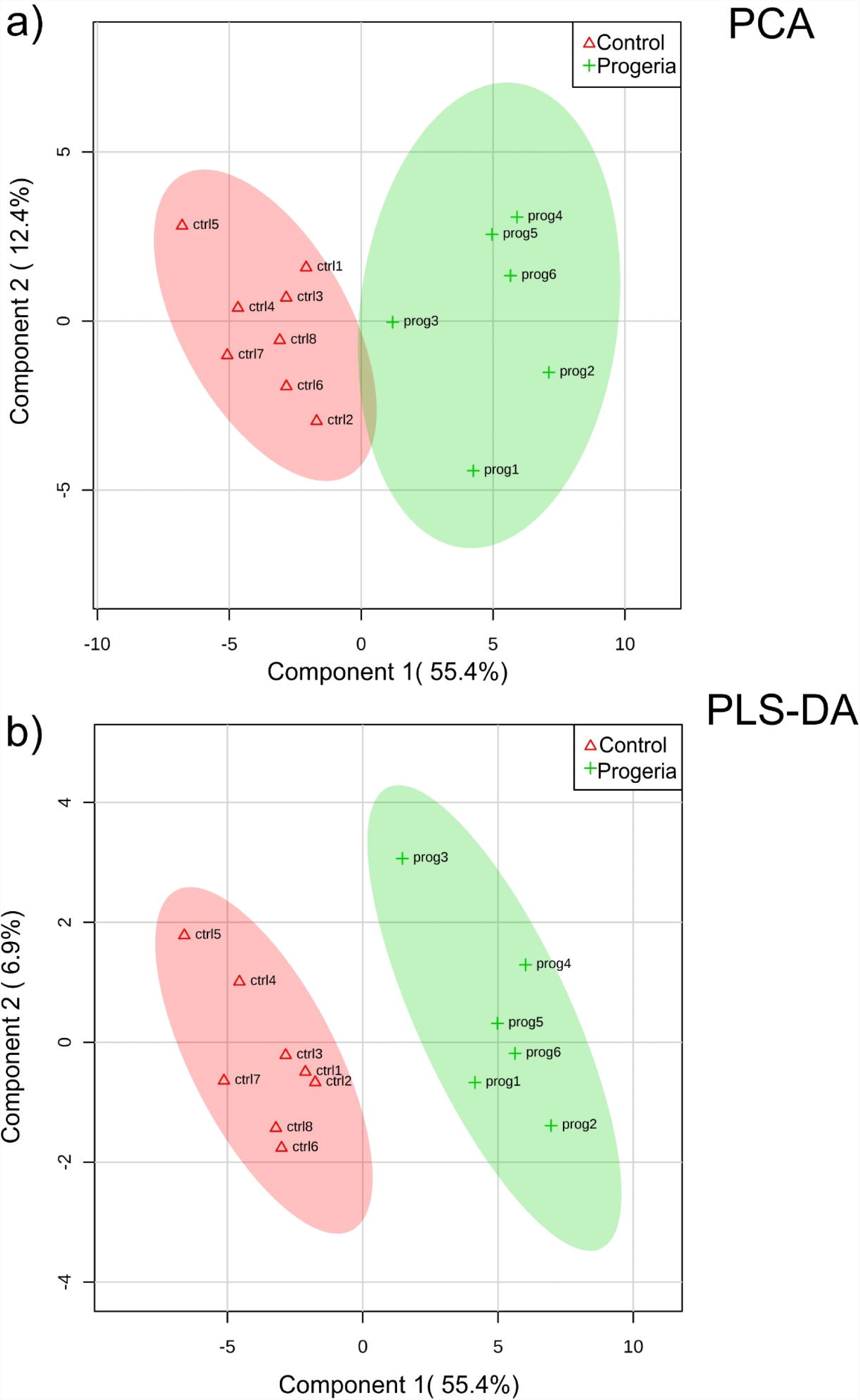
Multivariate analysis of the metabolomics data. a) Principal component analysis (PCA) 2D score plot and b) Partial Least Squares - Discriminant Analysis (PLS-DA) 2D score plot from the dataset with all of the features expected to be endogenous metabolites with statistical difference. The green (patients) and red (controls) areas represent the 95% confidence interval regions for each group.

In order to discover potential biomarkers for HGPS, ROC curves were constructed. The area under the ROC curve (AUC) is a well-described strategy for biomarker potential performance analysis, where the higher the AUC the more accurate is the model. As demonstrated in figure 2a, 6 AUC models were created, including different numbers of metabolites, varying from 2 to 40. The results demonstrate that the classification model using only two variables for the AUC resulted in a 0.803 value and 95% confidence interval (CI) ranging from 0.5∼1. Increasing the number of variables to 5 in the classification model, the AUC value increases to 0.912 and the 95% confidence interval ranges from 0.625∼1.

**Figure 2.**
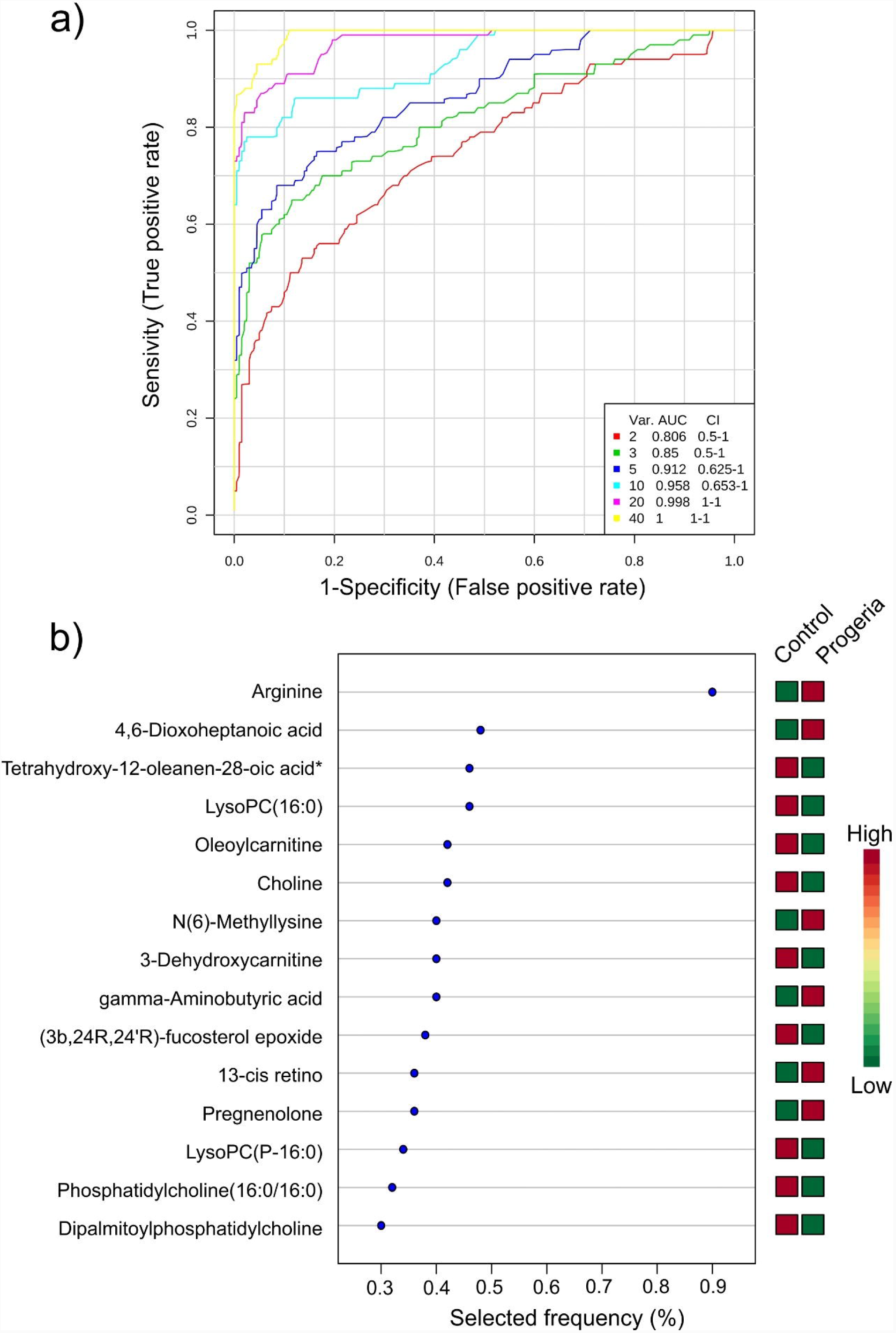
Potential biomarkers in diagnosing HGPS with metabolomics. a) Areas under the ROC curve (AUC) for different numbers of variables used to construct the ROC curves. The inset shows the number of variables used for the AUCs models. b) The most significant features of the ROC model downregulated (green) or upregulated (red) in patients with HGPS compared to controls.

Metabolites were ranked according to their capacity to distinguish between HGPS and Control subjects and the result is summarized in figure 2b. Metabolites are shown as either downregulated (green) or upregulated (red) in patients with HGPS. Furthermore, we performed classical univariate ROC curve analysis for individual biomarkers. Figure 3 shows the ranked metabolites based on area under ROC curve (AUROC), suggesting that both arginine and 5-hydroxytryptophol are robust upregulated candidates for HGPS biomarkers. Choline and phosphatidylcholine (16:0/16:0) on the other hand are robust downregulated candidates for HGPS biomarkers, showing potent diagnostic power. IAiming to validate the non-targeted metabolomic analysis, we performed targeted metabolomics based on LC-MS on a triple quadrupole using metabolite standards for prior calibration. Figure 4a shows ion chromatograms for arginine and ISTD. Each line represents one sample analyzed showing intensity and retention time in minutes. Figure 4b shows the calibration curve for arginine quantification. Our targeted method demonstrates an increase in arginine levels in samples from HGPS patients in the same manner as in the non-targeted approach as summarized in figure 4c. Data from arginine and other metabolites analyzed by triple quadrupole are summarized in table S3.

**Figure 3.**
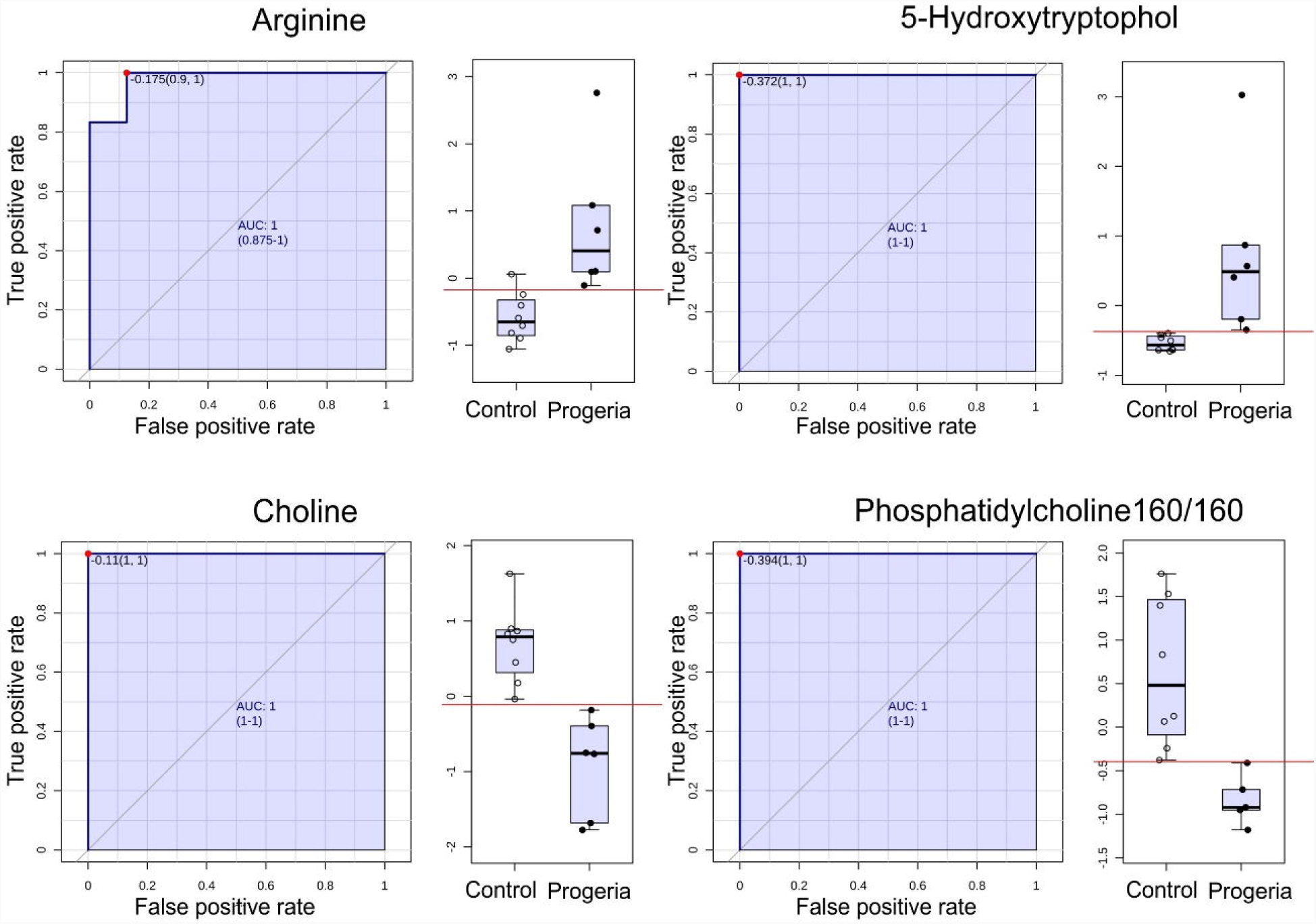
ROC curve analysis of individual biomarkers for HGPS based metabolomics. Top ranked metabolites based on area under ROC curve (AUROC) identified by the non-targeted metabolomics analysis of HGPS plasma samples. The rectangle to the right of the ROC curves show the individual values determined for each metabolite in the control and HGPS groups. The red line represents the mean value for the control group + two standard deviations.

**Figure 4.**
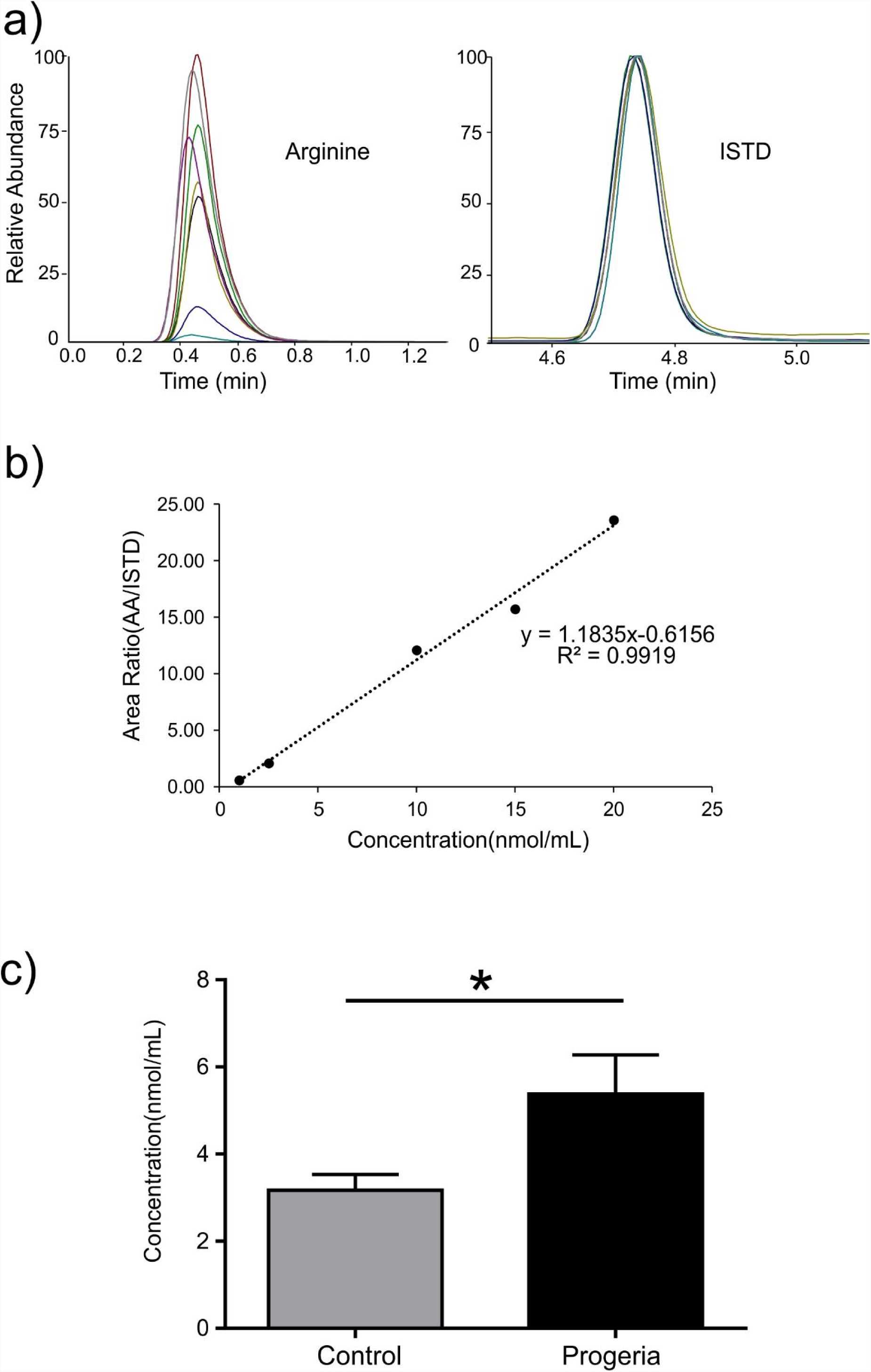
Targeted metabolomics performed by LC-MS/MS for biomarker validation. a) Extracted ion chromatograms for arginine and internal standard (ISTD). Each colored line represents one sample analyzed showing intensity and retention time (RT) in minutes. b) Calibration curve for arginine quantification. c) Graph shows arginine quantification in each group and bars represent SEM. * indicates P < 0.05.

Aiming to evaluate the most relevant metabolic pathways altered in patients with HGPS, pathway analysis was applied. Figure 5 shows an overview of Pathway Analysis, using only annotated metabolites identified to be significantly altered by HGPS. Figure 5a highlights pathways related to upregulated metabolites, suggesting alterations in fatty acids metabolism, glucose metabolism, and mitochondrial function. Figure 5b shows the metabolic pathways related to the downregulated metabolites in HGPS patients, demonstrating alterations related to phospholipids, phenylacetate and phosphatidylcholine metabolism, among other alterations.

**Figure 5.**
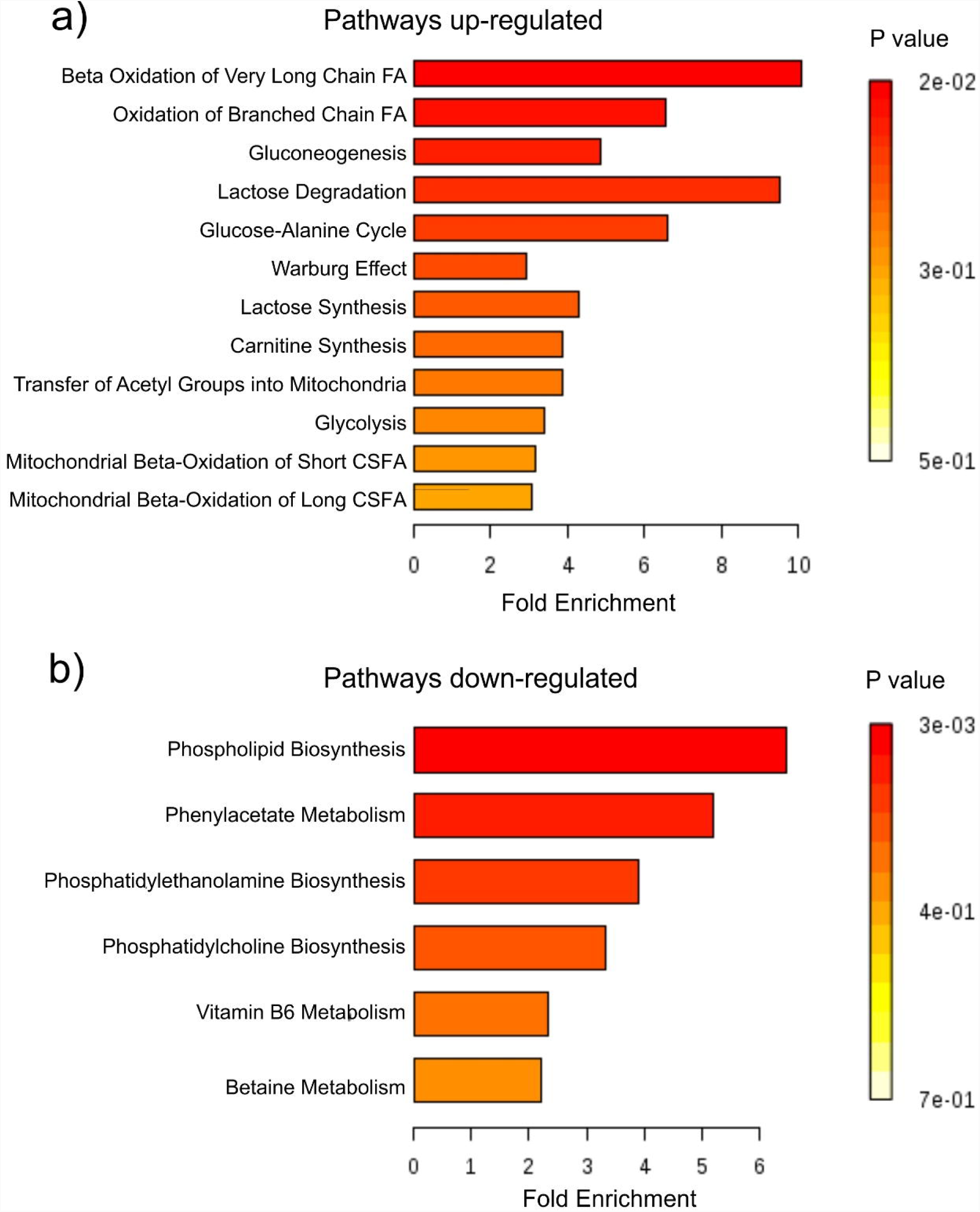
Metabolic pathways altered by HGPS. Overview of Pathway Analysis highlighting potential functional relationships between the set of annotated metabolites identified to be significantly altered by HGPS. a) Pathways related to upregulated and b) downregulated metabolites in the HGPS patients.

## Discussion

Aging is a complex biological process poorly understood at the molecular level. HGPS is a rare fatal disease where an extremely accelerated aging process is observed leading to premature death mainly related to heart complications. Despite great effort to increase knowledge about HGPS, biological biomarkers for this disease are not yet available, detailed disease mechanisms are still being investigated and there is currently no cure(*19*).

. In the present work, we created ROC curve models to identify metabolic HPGS biomarkers. As shown in results, biomarkers found to be statistically different between the HGPS and control groups are highly likely to be associated with premature aging based on performance in terms of both specificity and sensitivity. Furthermore, targeted analysis using high purity standards of the metabolites of interest previously identified in the non-targeted experiments are in accordance with the non-targeted strategies, showing similar results in terms of statistical significance.

A number of investigations used this methodology to study the mechanisms underlying the aging progression, and whether strategies such as exercise training and hormonal treatment can revert the metabolic changes induced by aging. In a recent study by Houtkooper et al, metabolomic hallmarks of aging were demonstrated, including affected pathways in both liver and muscle tissues, indicating a significant modification in fatty acid metabolism(*20*). In the present study, we found several compounds up or downregulated in the plasma of HGPS patients, highlighting a profound metabolic alteration compared to our control cohort. Aging metabolomic studies showed an increase in lactate and glucose suggesting changes in glucose/pyruvate and glycogen metabolism(*20*), in accordance with our data using HGPS plasma, where we observed an increase in glucose and lactide, a dimer of lactic acid (Table 1). Metabolomic studies in diabetic patients also demonstrate glucose and lactate increase(*21*). In addition to the glucose/pyruvate pathway alteration, we observed an increase in a long chain carnitine family molecule, Acetyl-L-Carnitine, associated with fatty oxidation. Interestingly, children in early-stage type 1 diabetes present elevated Acyl-Carnitine. Adult patients with type 2 diabetes may also present dysregulation of fatty acid oxidation, characterized by glucolipotoxicity(*22*). In this context, it is interesting to note that a recent study demonstrated that met-formin, a popular anti-diabetic biguanide, alleviates the nuclear defects and premature aging phenotypes in HGPS fibroblasts, perhaps constituting a promising therapeutic approach for life extension in HGPS(*23*). Furthermore, insulin resistance has been described in children with HGPS(*24*).

Mitochondria play a key role in several metabolic inborn errors as well as in the aging process, highlighting a decline in mitochondrial respiration(*20, 25, 26*). In this regard, carnitine metabolites are important during fatty acid oxidation in the mitochondria. We found an 11 fold increase in Acetyl-L-Carnitine and also in L-carnitine in HGPS patients reflecting a broad dysfunction in β-oxidation, indicating a diminished lipid transport capacity in the mitochondria(*27, 28*). On the other hand, we found some carnitine metabolites decreased in the plasma of HGPS, such as Decanoylcarnitine. Interestingly, fetal congenital disorders are associated with decreases in some carnitines, such as Decanoylcarnitine among others(*29, 30*). Collectively, these findings highlight the multiplicity of perturbations in lipid metabolism related to mitochondrial dys-functions in HGPS. These metabolic alterations may be related to the growth abnormalities observed in HGPS children(*31*).

During aging as well as in systemic metabolic dysfunction, amino-acid metabolism is significantly modified(*32, 33*). Previous publications demonstrate that branched-chain amino acids (BCAA) as well as methionine content in the diet changes mice lifespan. In the present work, we identified altered amino acid availability in HGPS patients’ plasma that was further investigated by targeted metabolomics. Our experiments showed a decrease in the levels of methionine and histidine, but in contrast levels of arginine and cystine were increased. Interestingly, Cheng and coworkers demonstrated that amino acids concentrations, such as Histidine, might be related to human longevity (*34*). Regarding BCAA no changes were observed, as well as in other important amino-acids such as proline and alanine. In agreement with our findings, Houtkooper et al showed that methionine is decreased in the plasma of aged mice and no changes were observed in BCAA(*20*). Interestingly, choline supplementation seems to improve cognitive function and is an important strategy to ameliorates Alzheimer’s disease pathology, a pathological process typically associated with advanced age(*35, 36*). Furthermore, choline converts homocysteine, a neurotoxic amino acid in methionine(*37, 38*), also found to be decreased in the HGPS patients in the present work.

Altered metabolic processes can lead to the formation of toxic metabolites as well as alterations in acid-base equilibrium. Interestingly, the 4,6-dioxoheptanoic acid, also known as Succinylacetone, a medium-chain keto acid, and derivative metabolite, was found in higher levels in HGPS plasma. Succinylacetone can rise due to abnormal activity of the enzyme fumarylacetoacetase, being suggested as an acidogenic, oncometabolite and a metabotoxin. Of note, aging and progeria course with a hypertrophic cardiac process, that dramatically increases the risk of severe cardiac complications. In this context, patients with hypertrophic cardiomyopathy are reported to have an increased level of this metabolite(*39, 40*).

Our study has limitations imposed by the cohort size used. As indicated in methods/results, we analyzed only 6 HGPS patients’ samples, a small number for a biomarker investigation and disease mechanism comprehension. However, HGPS is an extremely rare disease, as emphasized by the fact that in a 200 million people country like Brazil, only one donor was recruited. The 5 other samples from our cohort were donated by The Progeria Research Foundation which collects patients’ samples worldwide. These samples come from children with different genetic backgrounds, most probably contain different contaminants, were subject to distinct sample handling procedures and time of storage. Human genome databases show that the interindividual differences are very extensive between distinct populations. From the 40.000.000 variant polymorphic DNA sites predicted, some are rare and present only in a person or his family, ethnicity or country, which may reflect in their plasma metabolome(*41*). Remarkably, in view of the expected variability and the great possibility that the diverse genetic backgrounds might influence the metabolic plasma levels, our approach based in the multivariate analysis of multiple metabolites was capable to clearly separate patients from controls, generating an important biomarker profile related to the disease, even using a very small sample size.

In summary, the present work applied a powerful metabolomics pipeline based in liquid chromatography coupled to high-resolution mass spectrometry along with multivariate statistics and pathway analysis. We were able to identify putative circulating biomarkers for the disease that may be interesting targets for pharmacological treatment, nutritional supplementation and for diagnosing and follow up of HGPS patients.

## Conclusions

The present study reports for the first time a metabolic profiling with LC-MS based metabolomics of premature aging in patients with HGPS. We identified a total of 40 known metabolites differentially expressed between HGPS and age and sex-matched controls. Creating a panel with the most distinct metabolites, we identified circulating putative biomarkers candidates with high accuracy for group classification based on ROC curve models. Changed metabolic pathways involved fatty acids, amino-acids, and sphingolipids, among other metabolic pathways. Taken together these alterations impact the cellular energy use, enzyme activities, and cell signalling, creating new insights into the molecular mechanisms underlying premature aging associated with HGPS.

## Methods

### Sample preparation

Plasma samples were obtained from 5 HGPS patients kindly donated by The Progeria Research Foundation (www.progeriaresearch.org), including 3 females (2.3, 4.7, 12.2 years old) and 2 males (8.5, 10.2 years old). An additional sample was obtained from a Brazilian HGPS patient (female 8.4 years old) at the Federal University of Paraná as summarised in Table S1. The average age for HGPS patients was 7.7±1.4 years (mean±SD). As controls, we used 8 healthy donors of both genders (4 males and 4 females) with a mean age of 8.6±2.3 years (p= 0.4154).

### Ethics statement

Parents or the legal guardians of all controls and of the Brazilian patient have given full written informed consent for participation in the study. The study has been approved by the Ethics Committee of the Instituto Nacional de Cardiologia, number - 27044614.3.0000.5272 and the Department of Pediatrics from the University Hospital of the Federal University of Paraná. All procedures were in accordance with the ethical standards of the responsible local Ethics Committees and with the Helsinki Declaration of 1975, as revised in 2000.

### Metabolomics

Blood samples in Brazil were collected using EDTA tubes and plasma was obtained by centrifugation for 10 min at 10.000g (Megafuge 8R, Thermo Scientific, USA). The Progeria Research Foundation disposed frozen plasma samples. For metabolomic experiments, plasma proteins were precipitated with methanol (3:1 (v/v)) at −20°C for 1h. After protein precipitation, samples were centrifuged for 10 min at 14.000g, at 4°C, supernatants were collected and dried in a SpeedVac Concentrator (SPD111v, Thermo Scientific, USA). The metabolites were then reconstituted with a dilution factor of 3 in methanol/water (1:9 (v/v)). 5nM of deuterated testosterone (D3-Testerone, purchased from LGC Standards; London, England) was spiked and used as an internal standard (ISTD) for coefficient of variance (CV) calculation. For quality control (QC), a pool of all the analyzed samples was prepared.

### Non-targeted metabolomics

For liquid chromatography-tandem mass spectrometry (LC-MS) analysis, 5µl volumes of each sample were analyzed in triplicate. As a blank control, methanol/water followed the same steps and was used as background for data analysis as previously described(*15*). Between samples, a washing protocol was performed. Samples were analyzed in a random sequence and the QC sample was analyzed 5 different times along the experiment.

Samples were analyzed by Dionex Ultimate 3000 UHPLC coupled to a Q-Exactive Plus high-resolution mass spectrometer (Thermo Scientific, USA). LC separations were obtained using a 2.1 × 50mm ZORBAX 1.8µm C18 column (Agilent, USA). Mobile phases used were: phase A) water with 0.1% formic acid and 5mM ammonium formate, and phase B) methanol with 0.1% formic acid. Total run time was 30 min. The first 4 min of the run consisted of a linear gradient from 10% to 60% of B phase, followed by a 20 min linear gradient from 60% to 98% of B. After reaching 98%, a stable run with 98% of B was sustained for 3 minutes followed by 3 minutes of 10% of B solution to regenerate the column pumped at 450 μL/min with a column temperature of 55°C and the sample chamber held at 7°C acquired in positive mode. The data were obtained with the MS detector in full-scan mode (Full-MS) with the data-dependent acquisition (dd-MS2) for the top-10 most abundant ions per scan(*15*), with settings: In-source CID 0.0 eV, micro scans = 1, resolution = 70,000, AGC targeted 1e6, maximum IT = 50 ms, scan range 67 to 1000 m/z, spectrum data = Profile. Detector setting for dd-MS2 were: micro scans = 1, resolution = 17,500, AGC targeted 1e5, maximum IT = 100 ms, loop count = 10, isolation window 2.0 m/z, NCE 15, 35, 50, spectrum data = profile, underfill ratio = 1.5%, charge exclusion = unassigned, dynamic exclusion = 6s.

### Data analysis and statistics

Data were analyzed by Compound Discoverer 2.1 (Thermo Fischer, USA). For compound detection a mass tolerance of 5 ppm was accepted to extract ions with a minimum of 1.000.000 peak intensity; for compound consolidation, a 0.2 min of retention time tolerance was employed. The ChemSpider search including BioCyc and Human metabolome database (HMDB)(*16*) was used with 5ppm mass tolerance as well as the mzCloud search. In the non-targeted method, the identification of nom-novel metabolites was based on accurate mass and tandem mass spectra, without chemical standards references, providing a level 2 identification (putatively annotated). The samples were analyzed in triplicate. A principal component analysis (PCA) was performed to evaluate the experimental reproducibility and the QC samples were used to identify the reproducibility throughout experiments. Data of the triplicate injection experiments were unified and the average was used as a unique value. Data were scaled by auto-scaling. For statistical analysis, group area data from control vs patient data fold change was calculated and the p-value per group was calculated by t-test. Compounds that presented p<0.05 after adjustment using p-value (FDR) cutoff of 0.1 were considered statistically different. Chromatogram visualization and base peak chromatogram figure generation were performed using MZmine 2.26 software(*17*) and metabolomics statistics data was performed using MetaboAnalyst (*18*). The curated data matrix was used to generate a model for sample class discrimination via PCA and Partial Least Squares - Discriminant Analysis (PLS-DA) using online MetaboAnalyst (*18*). The model quality was analyzed by the goodness-of-fit parameter (R2) and the goodness-of-prediction parameter (Q2). For biomarker analysis, multivariate ROC curve based exploratory analysis was performed using a classification method (SVM) and feature ranking method (SVM built-in) applied to the statistically different metabolites between the two groups.

### Targeted metabolomics

Amino-acid mixture standard was purchased from Sigma-Aldrich (São Paulo, Brazil) and D3-testosterone (ISTD) from LGC Standards (London, England). Amino acid quantification was carried out using a TSQ Quantiva from Thermo Scientific (San Jose, USA) with a Dionex Ultimate 3000 HPLC system (Germering, Germany). Chromatographic separation was achieved using a reversed phase column (C18 Zorbax, 50 × 3 mm, 1,7 μm, Agilent, Santa Clara, USA). The analyte was eluted from the column using a gradient with the eluent changing from 5% to 100% methanol in water within 3 min. The column was washed for 1.2 min in 100% methanol and equilibrated for 3 min at the initial eluent composition. All solvents contained 0.1% formic acid. The flow rate, column temperature, and injection volume were 300 μL/min, 40°C and 5 μL, respectively.

Amino-acids were monitored by selected reaction monitoring (SRM) in the positive ion mode. The transitions selected for amino-acid quantification and ISTD are listed in Table S2. The curve was constructed using a mix of amino-acids in triplicate at 1; 2,5; 5; 10 and 20 nmol/mL. All samples were spiked with D3-Testosterone (ISTD) at 5 ng/mL. The area ratios of the total extracted ion of the product ions and the product ion of the IS were plotted versus the concentration.

### Statistical analysis

Data are presented as mean ± SEM. Two-tailed Student’s *t*-test was used. We did not use statistical methods to predetermine sample size; samples sizes were determined on the basis of sample availability. The non-targeted metabolomics statistics is described in detail with the metabolomics data analysis methods above. Values of P < 0.05 were considered statistically significant using GraphPad Prism 6.0 (GraphPad Software, USA).

## Acknowledgments

We are grateful to The Progeria Research Foundation for the availability of plasma samples, to Edna Aleixo from the Federal University of Rio de Janeiro for assistance with the importation process and to the Laboratório de Apoio ao Desenvolvimento Tecnológico (LADETEC) of the Institute of Chemistry of the Federal University of Rio de Janeiro for providing high quality infrastructure for the LC-MS analysis.

## Author contributions

GM and ACCC conceptualized the study and wrote the manuscript; GM, CGMS, JAME, GPCE, FCSN, GC, GBD, LM, ACCC acquired and analyzed the data; VOC, GBD, FCSN, ACCC critically revised the study and the manuscript.

