## supplementary material for "Metabolomic profiling revels systemic signatures of premature aging induced by Hutchinson-Gilford Progeria Syndrome"

**Supplementary Figure 1. Total ion chromatogram (TIC) of all analyzed samples**

a) TIC of Control plasma metabolomics. b) TIC of HGPS plasma metabolomics.

**Supplementary Figure 2. Evaluation of coefficient of variation in the non-targeted metabolomics methods**

**a)** Extracted ion chromatograms of the internal standard (ISTD) in the non-targeted method for the experimental samples and the Quality control (QC) samples. B) Calculation of the coefficient of variation (CV).

**Supplementary Figure 3. Partial Least Squares - Discriminant Analysis (PLS-DA) cross validation and permutation details**

a) Cross validation details of the PLS-DA. The asterisk represents the value selected. b) Permutation test performed using a set of 1000 permutation numbers.

Supplementary table 1. Gender and age from six patients and 8 controls from different families. All HGPS patients had the c1824C>T mutation.

| ID | Gender | Age at sample collection | Genotype |
| --- | --- | --- | --- |
| Patient 1 (L) | Female | 8.4 | LMNA Exon 11 c.1824C>T (p.Gly608Gly) |
| Patient 2 (191) | Female | 2.3 | LMNA Exon 11 c.1824C>T (p.Gly608Gly) |
| Patient 3 (113) | Female | 12.2 | LMNA Exon 11 c.1824C>T (p.Gly608Gly) |
| Patient 4 (194) | Male | 10.2 | LMNA Exon 11 c.1824C>T (p.Gly608Gly) |
| Patient 5 (009) | Male | 8.5 | LMNA Exon 11 c.1824C>T (p.Gly608Gly) |
| Patient 6 (092) | Female | 4.7 | LMNA Exon 11 c.1824C>T (p.Gly608Gly) |
| Control 1 | Female | 9.6 | - |
| Control 2 | Female | 7.0 | - |
| Control 3 | Male | 9.3 | - |
| Control 4 | Female | 9.5 | - |
| Control 5 | Male | 3.7 | - |
| Control 6 | Female | 10.5 | - |
| Control 7 | Male | 7.7 | - |
| Control 8 | Male | 11.9 | - |

Supplementary table 2. Transitions used in SRM experiment for quantification of metabolites

| Metabolites | Parent Mass  (m/z) | Product Mass  (m/z) |
| --- | --- | --- |
| Alanine | 90.2 | 44.2 |
| Arginine | 175.2 | 70.2 |
| Cysteine | 122.2 | 59.1 |
| ISTD-D3-Testosterone | 292.3 | 87.1 |
| Phenylamine | 166.1 | 30.3 |
| Glutamic Acid | 148.2 | 110.2 |
| Glutamine | 147.3 | 86.2 |
| Histidine | 156.3 | 84.2 |
| Leucine/Isoleucine | 132.1 | 133.1 |
| Lysine | 147.1 | 70.2 |
| Methionine | 150.2 | 60.2 |
| Proline | 116.1 | 91.1 |
| Tyrosine | 182.1 | 72.2 |
| Threonine | 182.1 | 120.1 |
| Tryptophan | 205.2 | 188.1 |
| Valine | 118.1 | 97.2 |

Supplementary table 3. Targeted metabolomics of plasma metabolites

| Compound | Control | HGPS | Fold Change | p-value |
| --- | --- | --- | --- | --- |
| Alanine | 22.23±2.9 | 14.87±5.2 | 0.68 | 0.21 |
| Arginine | 3.17±0.3 | 5.37±0.9 | 1.70 | 0.02 |
| Cysteine | 2.93±0.1 | 3.01±0.1 | 1.03 | <0.01 |
| Phenylamine | 32.51±2.3 | 25.76±3.0 | 0.79 | 0.09 |
| Glutamate | 0.95±0.2 | 2.002±0.5 | 2.09 | 0.07 |
| Glutamine | 6.302±0.8 | 4.42±1.4 | 0.70 | 0.23 |
| Histidine | 2.77±0.5 | 0.59±0.5 | 0.33 | 0.01 |
| Leucine | 80.54±9.2 | 71.99±17.8 | 0.89 | 0.65 |
| Lysine | 7.59±0.8 | 5.83±1.4 | 0.77 | 0.29 |
| Methionine | 5.67±0.4 | 4.26±0.5 | 0.75 | 0.05 |
| Proline | 26.26±6.5 | 14.15±2.5 | 0.54 | 0.13 |
| Tyrosine | 10.86±3.1 | 6.048±3.7 | 0.58 | 0.34 |
| Threonine | 10.66±3.1 | 5.825±3.8 | 0.58 | 0.34 |
| Tryptophan | 41.72±5.4 | 35.28±3.7 | 0.85 | 0.38 |
| Valine | 24.94±1.9 | 20.88±2.4 | 0.84 | 0.20 |
