## Supplementary figures and images for "Metabolomic profiling revels systemic signatures of premature aging induced by Hutchinson-Gilford Progeria Syndrome"

### sup. figure 1

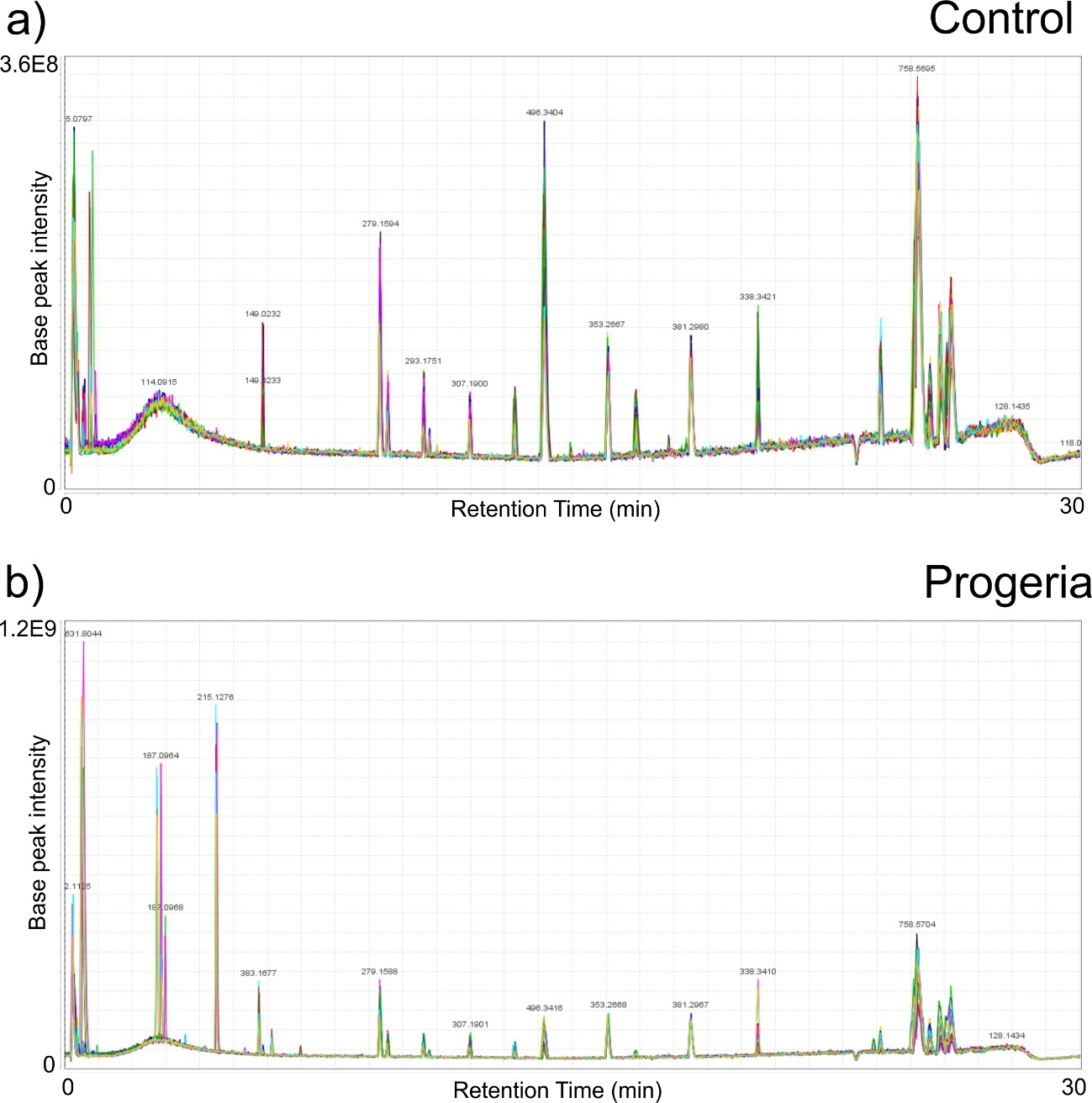

### sup. figure 2

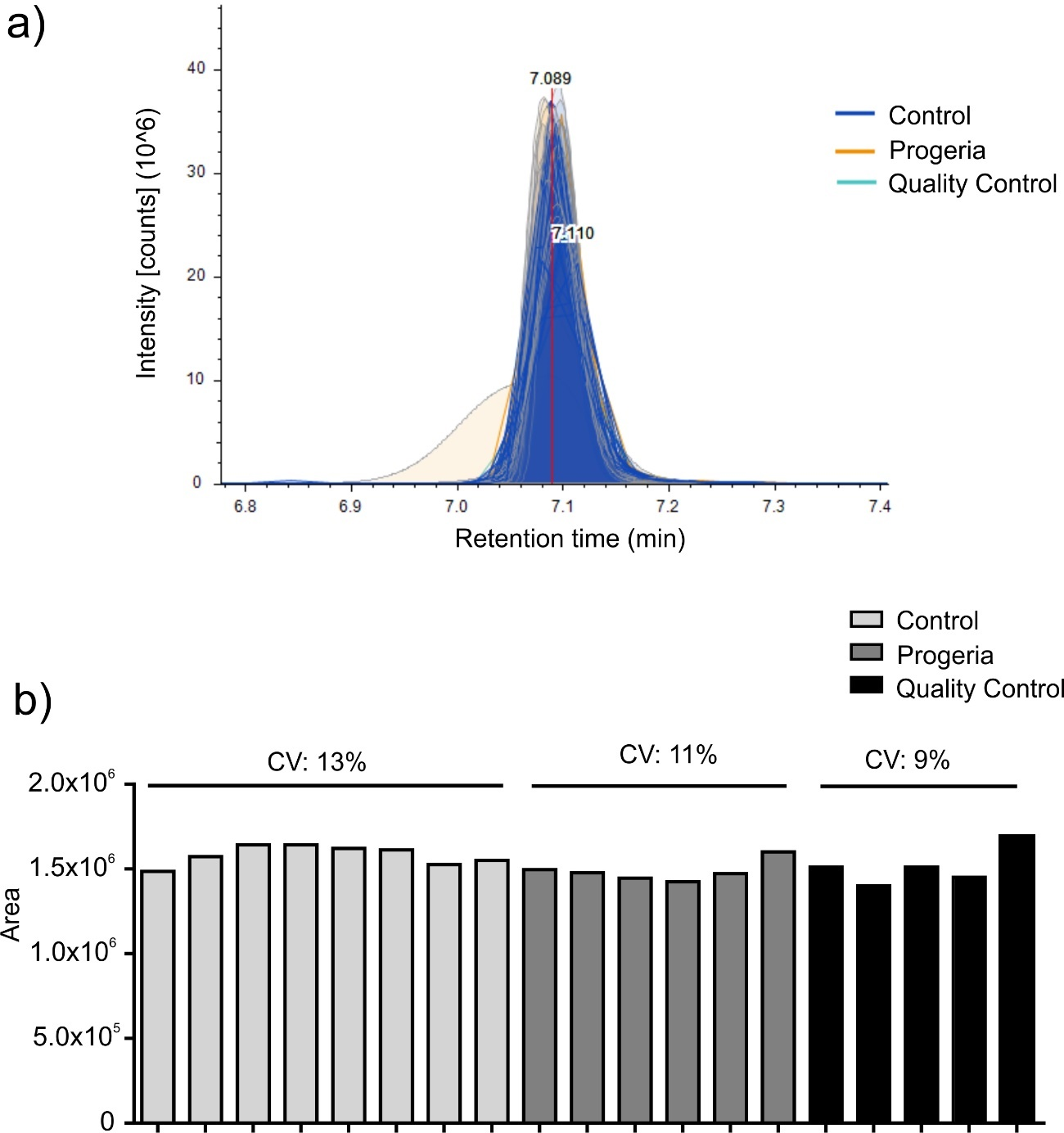

### sup. figure 3

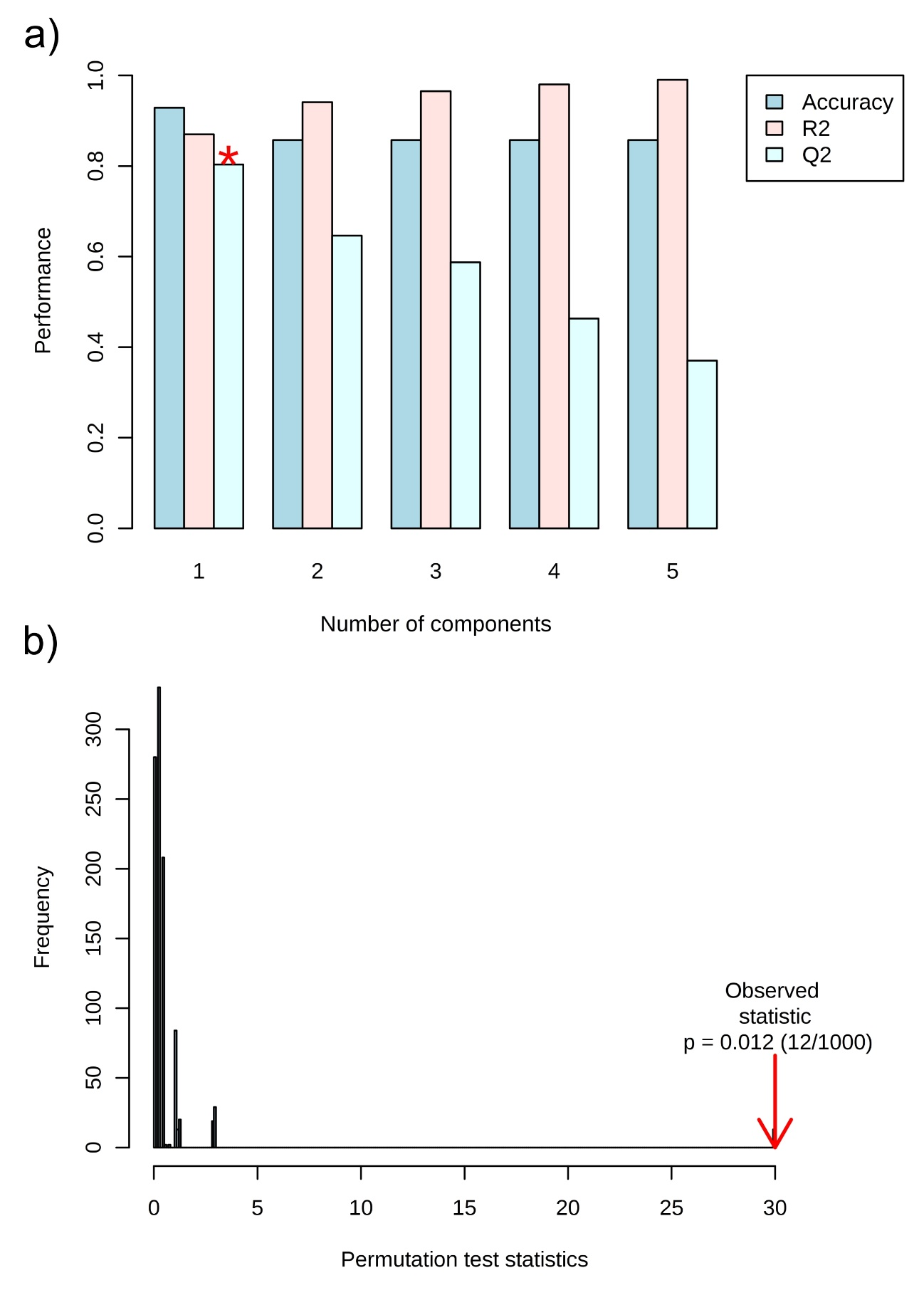
